# Evaluating the ability of spatial transcriptomics foundation models to learn multi-scale spatial variation

**DOI:** 10.64898/2026.08.01.742217

**Authors:** Dhanav Handa, Cristina Martin-Linares, Genevieve Stein-O’Brien, Jonathan Ling, Uthsav Chitra

## Abstract

Spatial gene expression results from the superposition of multiple sources of variation in gene expression across different spatial scales, including local microenvironment-associated variation and global spatial gradients. Spatial foundation models (SFMs) are large-scale machine learning models trained on cohorts of spatial transcriptomics (ST) data that, in principle, learn the different sources of spatial variation in gene expression. However, the embeddings learned by SFMs are difficult to interpret, and it remains unclear whether they fully capture such spatial variation. Here, we develop SAFFRON, a sparse autoencoder (SAE)-based framework for interpreting and evaluating SFMs. SAFFRON uses a Matryoshka SAE to decompose dense SFM embeddings into sparse, human-interpretable features and evaluates whether these features correlate with known sources of spatial variation. Using SAFFRON, we systematically benchmark the ability of several recent SFMs to identify local and global spatial variation in gene expression. We find that only two SFMs learn global spatial gradients more accurately than naive, non-foundation model baselines; further investigating one such SFM, SAFFRON reveals that these gradients are concentrated in a small subset of sparse, human-interpretable SAE features. On the other hand, *no* SFM learns local patterns (A*β* plaque microenvironment-associated patterns) more accurately than such baselines. Our findings suggest that current SFMs do not systematically learn multi-scale spatial variation in gene expression.

**Code:** SAFFRON is available at https://github.com/chitra-lab/SAFFRON.

## 1 Introduction

Spatial gene expression results from the superposition of multiple sources of spatial variation at different spatial scales. Such sources of spatial variation include localized microenvironments around proteins, cells, or tissues, e.g. pathological abnormalities around A*β* plaques in Alzheimer’s disease [1–4]; spatial gradients that modulate cellular differentiation and identity, e.g. in developing systems and the brain [5–8]; and the global organization of tissues into spatial domains, or discrete regions with distinct cell type composition and marker gene expression (e.g. the different layers of the cortex) [9–13].

Spatial transcriptomics (ST) technologies provide high-throughput gene expression profiling at thousands to millions of spatial locations in a two-dimensional (2-D) tissue slice, enabling the systematic study of spatial gene expression [14–18]. However, a major difficulty in analyzing ST data is the low sequence coverage (i.e. high data sparsity) and/or limited gene panels of current ST technologies. Sequencing-based ST technologies (e.g. 10x Genomics Visium/VisiumHD [19, 20], Stereo-seq [21], Slide-seq/Slide-tags [22–24]) measure a small number of mRNA transcripts per spatial location (roughly 500–5,000 unique transcripts per location) while imaging-based ST technologies (e.g. MERFISH [25], Nanostring CosMx [26], 10x Genomics Xenium [27]) measure the expression of a small number of genes in a cell, usually in the hundreds.

Spatial foundation models (SFMs) have recently emerged as a new paradigm for ST data analysis [28–31]. SFMs are large-scale machine learning models trained in an unsupervised manner on large collections of ST datasets (a process called *pretraining*). After pretraining, given a (potentially unseen) ST dataset, an SFM computes an *embedding* vector, or a dense numerical vector intended to summarize molecular state, for each cell/spatial location in the dataset. In principle, because SFMs are pretrained across diverse ST datasets, their embeddings should encode spatial gene expression patterns that recur across multiple ST datasets but are difficult to detect in an individual ST dataset due to large data sparsity or limited gene panels—analogous to how the embeddings of modern large language models (LLMs) encode recurrent patterns of language use and meaning learned from large text corpora [32–34]. However, the embeddings learned by SFMs are difficult to interpret and are a “black box”. Nearly all evaluations of SFMs focus on clustering embeddings into discrete spatial domains [28, 29], and it remains unknown whether SFM embeddings capture the other, more subtle sources of spatial variation in gene expression—including localized microenvironments and global spatial gradients—as well as how to infer such multi-scale spatial variation from SFM embeddings.

Sparse autoencoders (SAEs) are a powerful framework for interpreting the embedding vectors learned by foundation models [35–37]. SAEs decompose each dense embedding vector into a *sparse* linear combination of learned feature vectors. By imposing sparsity, SAEs encourage individual learned features to capture distinct and (ideally) human-interpretable dimensions of information that are encoded in foundation model embeddings. SAEs have been widely used for interpreting semantic concepts learned by foundation models in natural language processing and computer vision [36, 38, 39], and for identifying biological processes learned by protein language models [40–42] and single cell transcriptomics foundation models [43–45]. In principle, applying SAEs to SFM embeddings may enable the discovery of spatial gene expression patterns that are captured by the SFM but obscured in the original SFM embeddings. However, there are two key challenges in adapting SAEs to SFMs: (1) selecting an SAE architecture suited to SFM embeddings, as spatial variation in gene expression is driven by biological processes that occur at multiple spatial scales; and (2) interpreting the features learned by SAEs, which requires associating such features with distinct sources of spatial variation.

In this work, we introduce a Sparse Autoencoder Framework For Representing Omics Natural variation (SAFFRON). SAFFRON is a mechanistic interpretability framework for identifying and interpreting the spatial variation learned by SFMs. SAFFRON implements a Matryoshka SAE [46], an SAE architecture designed to learn hierarchical features, to decompose SFM embeddings into interpretable features that describe spatial variation at different spatial scales. SAFFRON then evaluates whether the SAE-learned features are correlated with known sources of spatial variation, and uses orthogonal matching pursuit (OMP) [47, 48] to identify a minimal set of SAE-learned features that reconstruct such spatial variation. We apply SAFFRON to systematically evaluate 6 recent SFMs/single cell FMs across 8 different spatial datasets with known sources of spatial variation, including global spatial gradients and local microenvironment-associated expression patterns. We find that only two SFMs learn global spatial gradients more accurately than naive matrix factorization baselines that only use the observed gene expression data (i.e. no pretraining); further, by investigating the SAFFRON-learned features for one such SFM, we find that only *two* SAE-learned features are required to accurately reconstruct spatial gradients. However, *no* SFM learn local gene expression patterns (A*β* plaque microenvironment-associated patterns) more accurately than naive baselines. Our findings suggest that current SFM architectures do not systematically learn multi-scale spatial patterns of gene expression.

## 2 Methods

### 2.1 Spatial transcriptomics and foundation models

*Spatial transcriptomics* (ST) technologies measure RNA transcripts across different spatial locations (spots) in a 2-D tissue section [49]. For a given tissue section *T* ⊆ ℝ^2^, ST technologies measure: (1) a gene expression matrix **A** = [**a**_*i*_] ∈ ℝ^*N*×*G*^, where each row **a**_*i*_ = (*a*_*i*1_, …, *a*_*iG*_)^⊺^ ∈ ℝ^*G*^ is the gene expression vector for spot *i* = 1, …, *N* and the individual entries *a*_*ig*_ are the expression of gene *g* in the spot *i*; and (2) the spatial location matrix **S** = [**s**_*i*_] ∈ ℝ^*N*×2^ where **s**_*i*_ = (*x*_*i*_, *y*_*i*_) ∈ *T* is the spatial location of spot *i*. In imaging-based technologies, each spot *i* = 1, …, *N* typically represents an individual cell, while in sequencing-based technologies, a spot may be made up of multiple cells (e.g. in 10x Genomics Visium [19], each spot consists of 5-20 cells). We assume the tissue *T* = *R*_1_ ∪··· ∪ *R*_*P*_ is divided into *P* disjoint regions, called *spatial domains*, where each region has a different gene expression pattern.

*Spatial foundation models (SFMs)* are large transformer-based models—pretrained on a large volume of ST data—that learn “general purpose” representations **z**_*i*_ for each cell/spot *i* in a tissue slice. Specifically, given ST data (**A, S**), a foundation model *f* learns a *D*-dimensional vector **z**_*i*_ = *f* (**a**_*i*_, **s**_*i*_) ∈ ℝ^*D*^ for each spot *i*. The vector **z**_*i*_ is called the *embedding vector* for cell *i*. Typically, the embedding dimension *D* ≪ *G* is much smaller than the number of genes *G*.

Some foundation models only use single cell transcriptomics data (single cell RNA sequencing, or scRNA-seq), which consists of a gene expression matrix **A** but not the spatial locations **S** of cells. These foundation models *f*, which we call single cell foundation models, learn a low-dimensional embedding **z**_*i*_ only from a gene expression vector, i.e. **z**_*i*_ = *f* (**a**_*i*_).

### 2.2 Sparse autoencoders

Sparse autoencoders (SAEs) are a recent approach for deriving human-interpretable features from foundation model embeddings **z**, which are dense vectors whose individual dimensions are typically not interpretable [35]; see Figure S1 for examples from SFMs. Given foundation model embeddings **z**_1_, …, **z**_*N*_, an SAE consists of two components: (1) an *encoder* that maps each embedding vector **z**_*i*_ to a high-dimensional, sparse vector **h**_*i*_ = [*h*_*im*_] ∈ ℝ^*M*^ and (2) a *decoder* that aims to reconstruct the embedding vector **z**_*i*_ from the sparse vector **h**_*i*_:

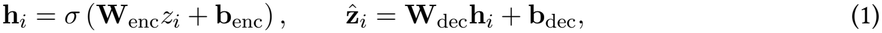

where **W**_enc_ ∈ ℝ^*M*×*D*^, **W**_dec_ ∈ ℝ^*D*×*M*^, **b**_enc_ ∈ ℝ^*M*^, **b**_dec_ ∈ ℝ^*D*^ are learnable parameters; *σ*(·) is a non-linear function; and 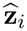 is a reconstructed version of the foundation model embedding vector **z**_***i***_.

We call the vector **h**_*i*_ the *SAE feature vector i*. for spot Typically the number *M* of features (i.e. the dimension of the vector **h**_*i*_) is larger than the dimension *D* of the foundation model embedding **z**_*i*_, i.e. *M > D*, as one aims to transform the dense vector **z**_*i*_ into a sparser, more interpretable vector **h**_*i*_. We call the vector ***η***_*m*_ = [*h*_*im*_] ∈ ℝ^*N*^, i.e. the *m*-th dimension of each feature vector **h**_*i*_ over all spots *i*, the *m-th SAE feature*, and we call its entries *h*_*im*_ the *activations* of the *m*-th SAE feature.

The SAE feature vectors **h**_*i*_ are learned by minimizing the loss function 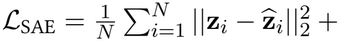 ℛ(**h**_*i*_), where ℛ(**h**_*i*_) is a regularization term on the vector **h**_*i*_. The sparsity of the SAE feature vector **h**_*i*_ is typically encouraged with either (1) a sparsity-inducing regularization term (e.g. the *l*_1_ regularization term ℛ(**h**_*i*_) = *λ·* ||**h**_*i*_||_1_ for some hyperparameter *λ >* 0) or (2) a “top-*K*” activation function *σ* = *σ*_*K*_(**v**) in (1), which sets *σ*_*K*_(**v**)_*i*_ = 0 if entry *i* is not in the top *K* entries of **v**, for some *K >* 0 [35, 37].

#### Matryoshka SAEs

Matryoshka SAEs are a recently developed extension of SAEs that aim to learn “nested” SAE feature vectors [46]. Early SAE features (i.e. ***η***_*m*_ for small *m*) should correspond to “coarse-grained” features while later dimensions (i.e. ***η***_*m*_ for large *m*) should correspond to “fine-grained” features. This nested structure is learned in the following manner. Let 1 ≤ *M*_1_ *< M*_2_ *<·*·· *< M*_*R*_ = *M* be a sequence of *R* indices, which we call prefixes. For each prefix *M*_*r*_, the Matryoshka SAE aims to reconstruct the SFM embedding **z**_*i*_ using only the first *M* entries of the feature vector **h**_*i*_; i.e. 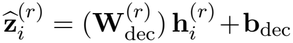, where 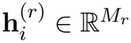 is a vector consisting of the first *M*_*r*_ entries of the vector **h**_*i*_ and 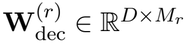 is a matrix consisting of the first *M*_*r*_ columns of the decoder matrix **W**_dec_.

The Matryoshka SAE loss 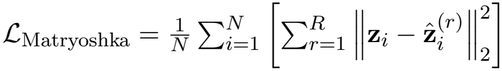 is the mean of the reconstruction losses across all prefixes *r* = 1, …, *R* [46]. The Matryoshka loss requires early entries of the SAE feature vector (i.e. the vectors **h**^(*r*)^ for small *r*) to reconstruct the entire foundation model embedding **z**, and thus ideally these early entries should capture the dominant sources of variation in gene expression; in contrast, later entries of the SAE feature vector should ideally concentrate on more fine-grained variation.

### 2.3 SAFFRON

SAFFRON is a mechanistic interpretability framework for identifying and interpreting the sources of spatial variation in gene expression encoded in SFM embeddings **z**. SAFFRON consists of two components.

First, SAFFRON implements a Matryoshka SAE with BatchTopK [50] sparse activation to learn SAE feature vectors **h**_*i*_ for each cell. We selected the SAE architecture via a systematic ablation on the STARmap brain dataset (Section 3.1) where we compared vanilla and Matryoshka SAE variants, sparsity constraints, and different hyperparameters *M, K*. See Appendix A for more details.

Second, SAFFRON evaluates how well the SAE features correlate with a pre-defined measure ***τ*** = [*τ*_*i*_] ∈ ℝ^*N*^ of spatial variation in gene expression, represented as a scalar value *τ*_*i*_ for each cell *i* = 1, …, *N* (see Section 3.1 for specific examples). We quantify such correlation by computing the maximum Spearman correlation *ρ* between the spatial measure ***τ*** and each SAE feature ***η***_*m*_ for dimension *m* = 1, …, *M*. For SFMs where the Spearman correlation *ρ* is large, we use orthogonal matching pursuit (OMP) [48] to identify a *minimal* set *S* ⊆ {1, …, *M*} of SAE features that linearly reconstruct the spatial variation measure ***τ***, i.e. ***τ*** ≈ ∑_*m*∈*S*_ *β*_*m*_***η***_*m*_ for some *β*_*m*_ ∈ ℝ. See [47, 51] for more details on OMP.

## 3 Results

### 3.1 Foundation models, datasets, and spatial patterns

We use SAFFRON to evaluate the embeddings of six transcriptomic foundation models (Table S1). These include four SFMs: Nicheformer [30] and scGPT-Spatial [29], which are transformer-based SFMs; Novae [28], a graph-based SFM; and SpatialFusion [31], a multimodal FM which integrates ST data with Hematoxylin and Eosin (H&E) images. We use three variants of Novae: *train-from-scratch* (TFS), where Novae is trained only on the target dataset; *fine-tune* (FT), where Novae’s pretrained weights are further trained on the target dataset; and *zero-shot* (ZS), where the pre-trained Novae is directly applied. For SpatialFusion, we evaluate four configurations: SpatialFusion-RNA (SF-RNA), which only uses the ST data; SpatialFusion-H&E (SF-H&E), which only uses the H&E image (when available); SpatialFusion-Both (SF-Both), which uses both ST and H&E; and a fine-tuned variant of SF-Both (SF-Both-FT). We do not fine-tune Nicheformer or scGPT-spatial as these methods do not have unsupervised fine-tuning capabilities. We evaluate two single-cell (i.e. non-spatial) foundation model baselines, Geneformer [52] and scGPT [53]. To run each spatial or single-cell foundation model, we follow the provided tutorials on GitHub. As non-foundation model baselines, we compare against PCA and SAFFRON’s SAE applied directly to the gene expression matrix **A**. Additional model-specific implementation and fine-tuning details are provided in Appendices B and C.

We evaluate these models using eight ST datasets which span multiple tissues and ST technologies and which exhibit known, quantifiable spatial patterns of gene expression (Table 1). The first six datasets in Table 1 exhibit spatial gradients of gene expression along a one-dimensional (1-D) spatial axis (e.g. in the cortex, genes exhibit expression gradients along the cortical depth axis), which we quantify using either GASTON [8], an algorithm which learns the 1-D axis of maximum spatial variation in a tissue (called the *isodepth*), or an author-provided 1-D coordinate (*pseudospace*) for the GI tract dataset [54]. Since both GASTON and the GI tract pseudospace coordinate depend on gene expression, we also evaluated the GI tract dataset using a morphological coordinate learned by MorphoGAM [55], which is computed only from cell positions (see Appendix Figure S2). The latter two ST datasets in Table 1 are brains with Alzheimer’s disease which exhibit local microenvironment-associated expression patterns around A*β* plaques (e.g. microglial accumulation). We use three distinct, cell-level metrics to quantify such local variation: (1) the disease-associated microglia (DAM) score *τ* ^(1)^, or the mean expression of the 26 DAM signature genes found by [56]; (2) the local microglial density *τ* ^(2)^, or the number of microglial cells within a 100 *µ*m radius of a cell defined by [57]; and (3) the distance *τ* ^(3)^ to the nearest plaque [57]. The latter two metrics are slightly correlated (median Spearman correlation *ρ* = −0.17 and *ρ* = −0.31 in each dataset).

**Table 1.** ST datasets analyzed in this work. Observations denote spots (Visium), bins (VisiumHD), or cells (otherwise)

| **Tissue** | **Species** | **Technology** | **# Observations** | **# Genes** | **Spatial variation** |
|---|---|---|---|---|---|
| Colorectal tumor [58] | Human | Visium | 3,900 | 11,008 | 1-D tumor-stroma axis |
| Kidney [27] | Mouse | VisiumHD | 106,302 | 3,856 | 1-D radial axis |
| Cerebral cortex [59] | Human | MERFISH | 17,450 | 282 | 1-D cortical depth axis |
| Cerebral cortex [59] | Human | MERFISH | 12,861 | 287 | 1-D cortical depth axis |
| GI tract [54] | Mouse | MERFISH | 2,060,051 | 1,815 | 1-D crypt-villus axis |
| Colon [20] | Human | Xenium | 260,232 | 226 | 1-D crypt-surface axis |
| Brain (Alzheimer’s disease) [1] | Mouse | STARmap | 72,165 | 2,766 | Local *Aβ* plaque microenvironment |
| Brain (Alzheimer’s disease) [57] | Mouse | MERFISH | 432,794 | 300 | Local *Aβ* plaque microenvironment |

### 3.2 SAFFRON evaluates the ability of SFMs to identify spatial gradients

We first use SAFFRON to evaluate whether SFMs learn spatial gradients of gene expression. For each dataset and model, we quantify gradient recovery as the maximum absolute Spearman correlation |*ρ*| between any single SAE feature ***η***_*m*_ learned by SAFFRON and a reference 1-D spatial axis *τ* (Section 3.1). Since the SAE feature ***η***_*m*_ is sparse by design, we calculate the correlation over all locations *i* within predefined spatial domains *R*_*p*_ obtained using GraphST [12] or GASTON [8]. We then average the maximum correlations across spatial domains to obtain a single score for each dataset and model. By taking the maximum correlation across SAE features, we assess whether any single SAE-learned feature captures the reference spatial gradient, which makes this metric intentionally favorable to SFMs. We further assess the robustness of this metric using permutation-based null comparisons and fixed SAE dictionary sizes (Figure S3).

We find (Figure 2a) that the SAE-learned features for Novae (either trained from scratch or finetuned) achieve the highest Spearman correlations (median |*ρ*| = 0.75 and 0.68, respectively), substantially exceeding the gene expression SAE baseline (median |*ρ*| = 0.38) and all other SFMs and single-cell foundation models (median |*ρ*| *<* 0.60). Noteworthily, SpatialFusion also exceeds the naive gene-expression SAE baseline, both from expression alone (median |*ρ*| = 0.59) and when it incorporates matched H&E histology (median |*ρ*| = 0.47, rising to 0.54 with fine-tuning), yet remains below Novae on all but one dataset. The remaining SFMs (scGPT-Spatial and Nicheformer) had Spearman correlations comparable to non-spatial foundation models and the naive SAE/PCA baselines, suggesting that these models are not able to adequately leverage spatial context during pre-training to capture continuous spatial gradients. For example, in a colorectal tumor region (Figure 2b, left), many genes exhibit gradients along a 1-D tumor core-to-tumor edge axis (Figure 2b, right, “Reference”) [8]. Using SAFFRON, we find that the Novae and SpatialFusion SAE features accurately identify such a 1-D tumor axis, while other SFMs learn spatially incoherent features that do not align with the known tumor axis (Figure 2b, right). Similarly, in the cerebral cortex, many genes and cell types exhibit smooth spatial gradients along the cortical depth axis (Fig. 2c, left; Fig. 2c, right, “Reference”). Only the Novae SAE features learn the cortical depth axis, while the other SFMs learned SAE features that are visually similar to the naive SAE baseline (Fig. 2c, right). We observe similar qualitative findings for the four other ST datasets we analyzed (Figure S4).

**Figure 1:**
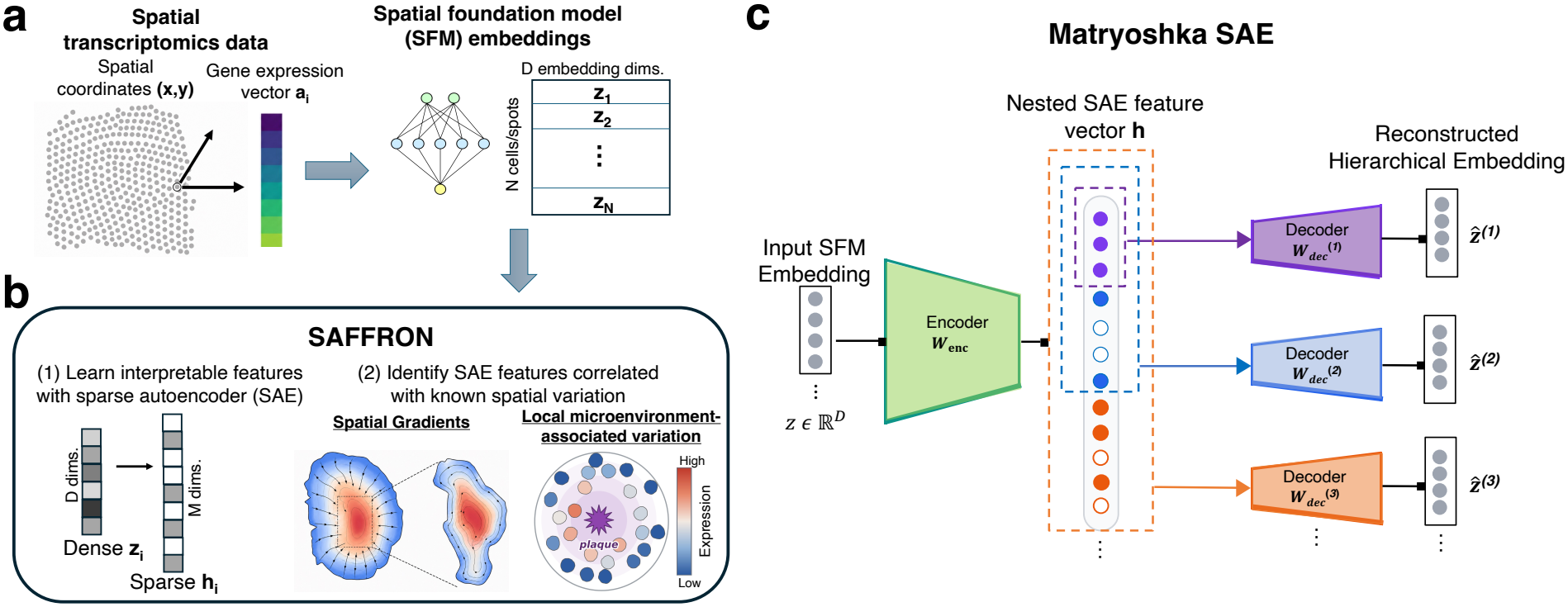
Schematic overview of SAFFRON. **(a)** (Left) Spatial transcriptomics (ST) data consists of measured gene expression vectors **a**_*i*_ at *N* different spatial locations (*x*_*i*_, *y*_*i*_) in a 2-D tissue slice. (Right) Spatial foundation models (SFMs) take in an ST dataset and learn embedding vectors **z**_1_, …, **z**_*N*_ for the *N* spatial locations. **(b)** SAFFRON uses a sparse autoencoder (SAE) to transform the dense embedding vectors **z**_*i*_ into sparse, human-interpretable vectors **h**_*i*_, called SAE feature vectors. SAFFRON then identifies which SAE features (i.e. dimensions of **h**) correlate with known spatial variation including continuous spatial gradients of gene expression and local microenvironment-associated variation in expression. **(c)** SAFFRON specifically uses a Matryoshka SAE architecture, which takes in an SFM embedding **z** and learns a “nested” vector **h** by reconstructing the SFM embedding **z** from different prefixes of **h**.

**Figure 2:**
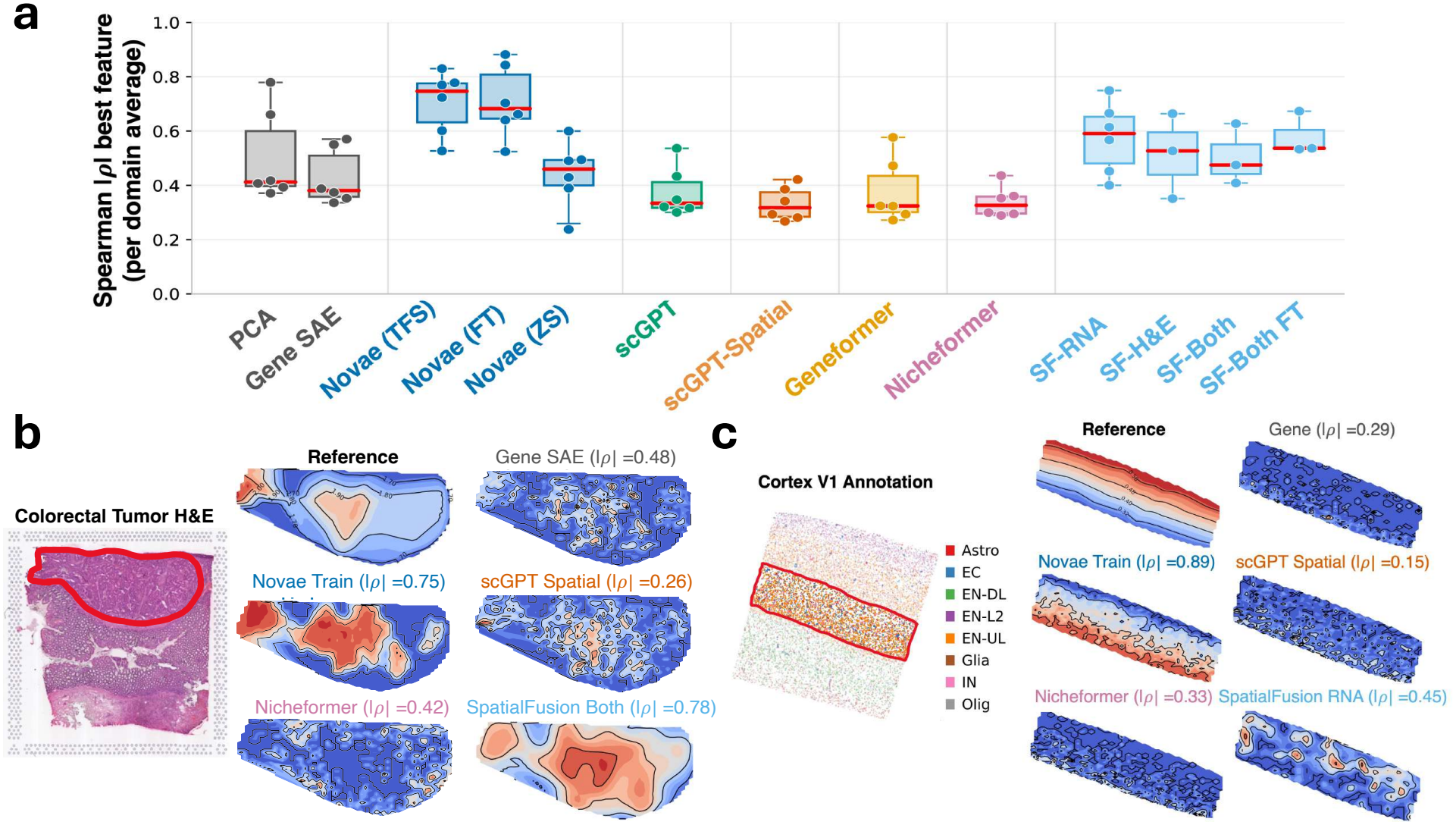
**(a)** Maximum absolute Spearman correlation |*ρ*| averaged across spatial domains *R*_*p*_ between a known reference 1-D spatial axis ***τ*** and SAFFRON SAE-learned features ***η*** derived from either normalized gene expression **a** or embeddings **z** of different SFM/single cell foundation models. **(b)** – **(c)** (Left) H&E for colorectal tumor (b) and cell type annotations for cerebral cortex (c). Circled region denotes spatial domain visualized on right. (Right) Reference 1-D spatial axis ***τ*** and SAFFRON SAE-learned feature with highest correlation to ***τ*** for select models shown in (a).

Overall, SAFFRON’s spatial interpretability framework reveals that Novae learns embeddings that encode continuous spatial gradients; that SpatialFusion also captures such gradients but less consistently across datasets; and that the other foundation models do not learn such gradients.

### 3.3 SAFFRON uncovers a small set of Novae SAE features that reconstruct spatial gradients

We next use SAFFRON to investigate whether Novae is able to reconstruct spatial gradients in an interpretable manner. While the preceding analysis (Section 3.2, Figure 2) showed that the Novae SAE features have relatively high correlation with known 1-D spatial axes, it is unclear whether these axes can be reconstructed by a small number of Novae SAE features—which could conceivably be done without any prior knowledge on the 1-D spatial axis—or whether accurate reconstruction depends on many features, and thus substantial prior knowledge of the 1-D axis is required.

We used SAFFRON’s orthogonal matching pursuit (OMP) approach to identify minimal sets of Novae SAE features that linearly reconstruct the reference 1-D tumor-to-stroma axis in a colorectal tumor sample (Figure 3a). We find (Figure 3b) that for both the Novae TFS and fine-tuned Novae models, only a small number of SAE features are needed to reconstruct the 1-D tumor axis. In particular, for Novae TFS, the 1-D axis constructed with *M* = 2 SAE features (Figure 3c) already achieves *R*^2^ *>* 0.8 with the reference 1-D axis. Interestingly, the zero-shot Novae also reconstructs the reference axis reasonably well using all *M* = 256 SAE features (*R*^2^ = 0.89, Figure 3b). The two Novae TFS SAE features selected by SAFFRON display spatial axes in different domains: one SAE feature captures the stroma-to-tumor-edge axis in the stromal domain (Figure 3d, left) while the other SAE feature captures the tumor edge-to-core axis in the tumor domain (Figure 3d, right).

**Figure 3:**
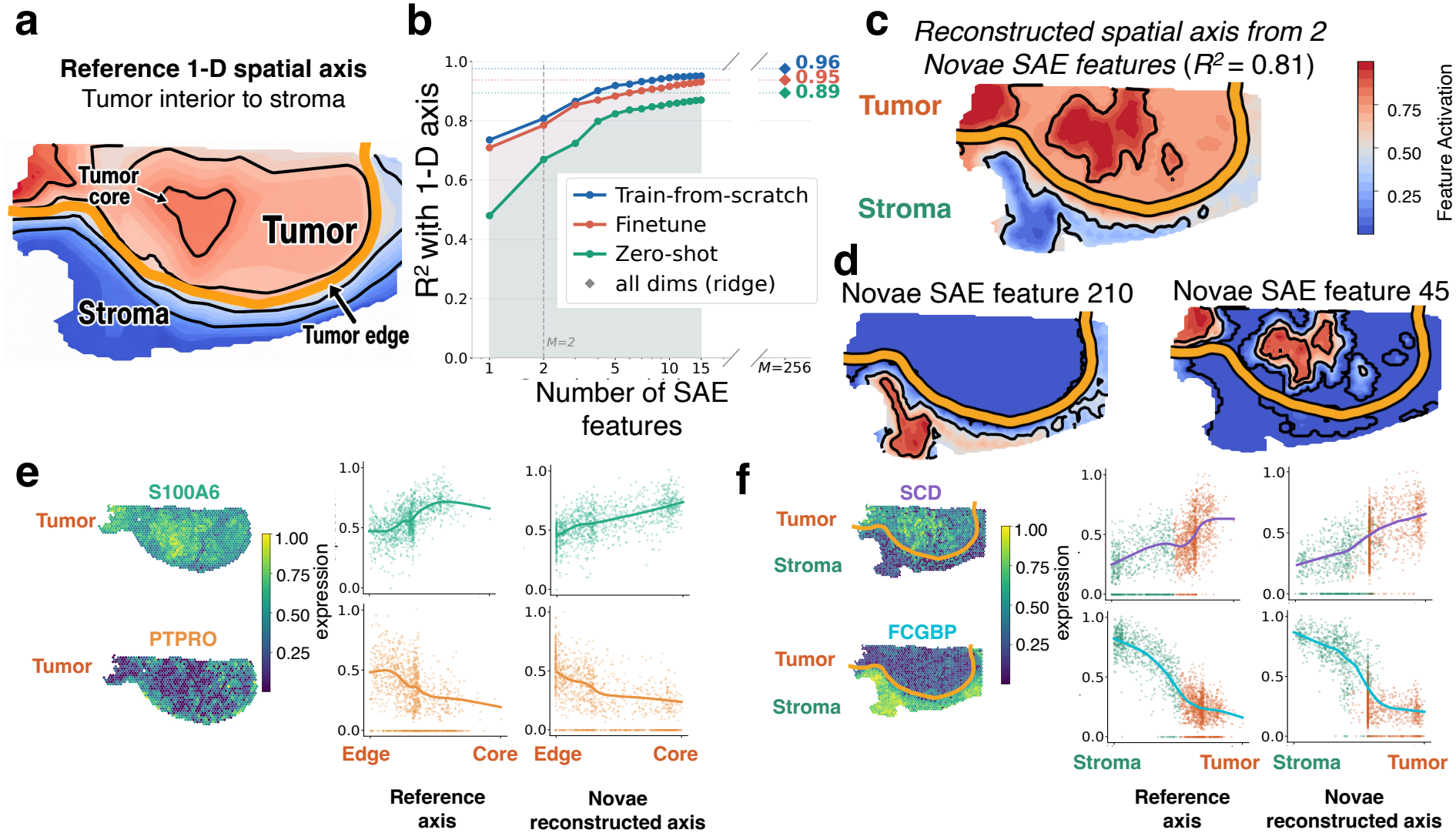
**(a)** Schematic of the reference 1-D spatial axis, oriented from tumor interior (core) to surrounding stroma, inferred by GASTON [8]. Amber line marks the inferred tumor–stroma boundary, and black lines are contours of equal 1-D coordinate. **(b)** Cross-validated *R*^2^ (coefficient of determination) of sparse OMP reconstruction of the reference 1-D spatial axis using SAE features from Novae TFS, fine-tuned, and zero-shot. **(c)** Spatial map of the *M* = 2 OMP reconstruction across the tumor interior–stroma boundary region. **(d)** Spatial visualizations of the two individual Novae SAE features ***η***_210_, ***η***_45_ selected by SAFFRON. **(e)** Within-tumor gene expression for S100A6 (top) and PTPRO (bottom), shown as spatial maps (left), versus the reference isodepth axis (center; x-axis: tumor edge to core), and versus the Novae reconstructed axis (right) **(f)** Across tumor-stroma gene expression for SCD (top) and FCGBP (bottom), shown as spatial maps with amber boundary contour (left), versus the reference axis (center; x-axis: stroma to tumor), and versus the Novae reconstructed axis (right).

The spatial tumor axis reconstructed by the Novae SAE features recapitulates spatial gene expression gradients both within the tumor region and across the tumor boundary. Within the tumor region, the two genes whose expression has the largest absolute Spearman correlation with the reference tumor edge-to-core axis are *S100A6* (correlation *ρ* = 0.59 with reference axis), which encodes a calcium-binding protein associated with tumor progression [60], and *PTPRO* (correlation *ρ* = −0.44 with reference axis), which encodes a receptor-type protein tyrosine phosphatase that suppresses colorectal cancer tumorigenesis and metastasis [61]. *S100A6* (resp. *PTPRO*) has an increasing (resp. decreasing) expression gradient from the tumor edge to the core measured with either the reference tumor axis or the reconstructed axis from the Novae SAE features (Figure 3e). Similarly, across the tumor-stroma boundary, two genes highly correlated with the reference tumor-to-stroma axis are *SCD*, a fatty-acid desaturase associated with colorectal cancer metastasis [62], and *FCGBP*, a tumor suppressor in colorectal cancer enriched in the adjacent stromal region [63]. These two genes have a similar expression gradient across the tumor-stroma region whether this gradient is measured using the reference tumor-stroma axis or with the Novae-reconstructed axis (Figure 3f). Together, our results demonstrate how, by utilizing Novae in conjunction with SAFFRON, we can identify minimal and human-interpretable subsets of Novae SAE features that capture biologically meaningful spatial gradients within the tumor and across the tumor boundary.

### 3.4 SAFFRON evaluates whether SFMs identify local A*β* plaque microenvironment in Alzheimer’s disease

We next used SAFFRON to evaluate whether SFMs capture localized variation in gene expression around A*β* plaques using two ST datasets from brains with Alzheimer’s disease [1, 57]. Alzheimer’s disease is marked by changes in cell type organization (e.g. increased microglia proportion) locally around A*β* plaques [1–4], but such changes are often difficult to identify from individual sparse ST datasets. We thus aimed to test whether SFMs can identify such local, microenvironment patterns, which we quantify using the DAM score, local microglial density, and distance to nearest plaque metrics (Section 3.1).

We find (Figure 4a) that for all three metrics, the naive gene expression baselines (PCA, SAE) identify features with similar (and sometimes larger) correlation to the metric than the SFMs. For example, for the DAM score metric, PCA and SAFFRON’s SAE on the raw gene expression matrix yield features with much larger Spearman correlation with the DAM score (median PCA |*ρ*| = 0.59; median SAE |*ρ*| = 0.58) than the SAE features learned by any SFM or single-cell FM (Figure 4b, top). Interestingly, SAFFRON finds that the SAE features for Geneformer, a non-spatial foundation model, are more correlated with the DAM score than the SAE features of the spatial foundation models. Further, the SAE features for all methods have low correlation to the local microglial density metric (median |*ρ*| = 0.16–0.27), with no clear advantage for the foundation models. Finally, for the distance to nearest plaque metric, we found that applying SAFFRON’s SAE to the Novae embeddings yields a SAE feature with somewhat larger correlation than the naive PCA/SAE baselines (Novae finetuned |*ρ*| = 0.33; PCA |*ρ*| = 0.23; Gene SAE |*ρ*| = 0.25), while SpatialFusion is comparable to the baselines (|*ρ*| = 0.26) and the remaining SFMs fall below them. However, by visualizing the most correlated foundation model SAE feature (SF-RNA) in a representative region, we qualitatively observe that this feature does not identify the distance to plaque metric (Figure 4b, bottom). Our results suggest that SFMs do not substantially improve over simpler baselines in recovering local A*β* plaque-associated spatial variation.

**Figure 4:**
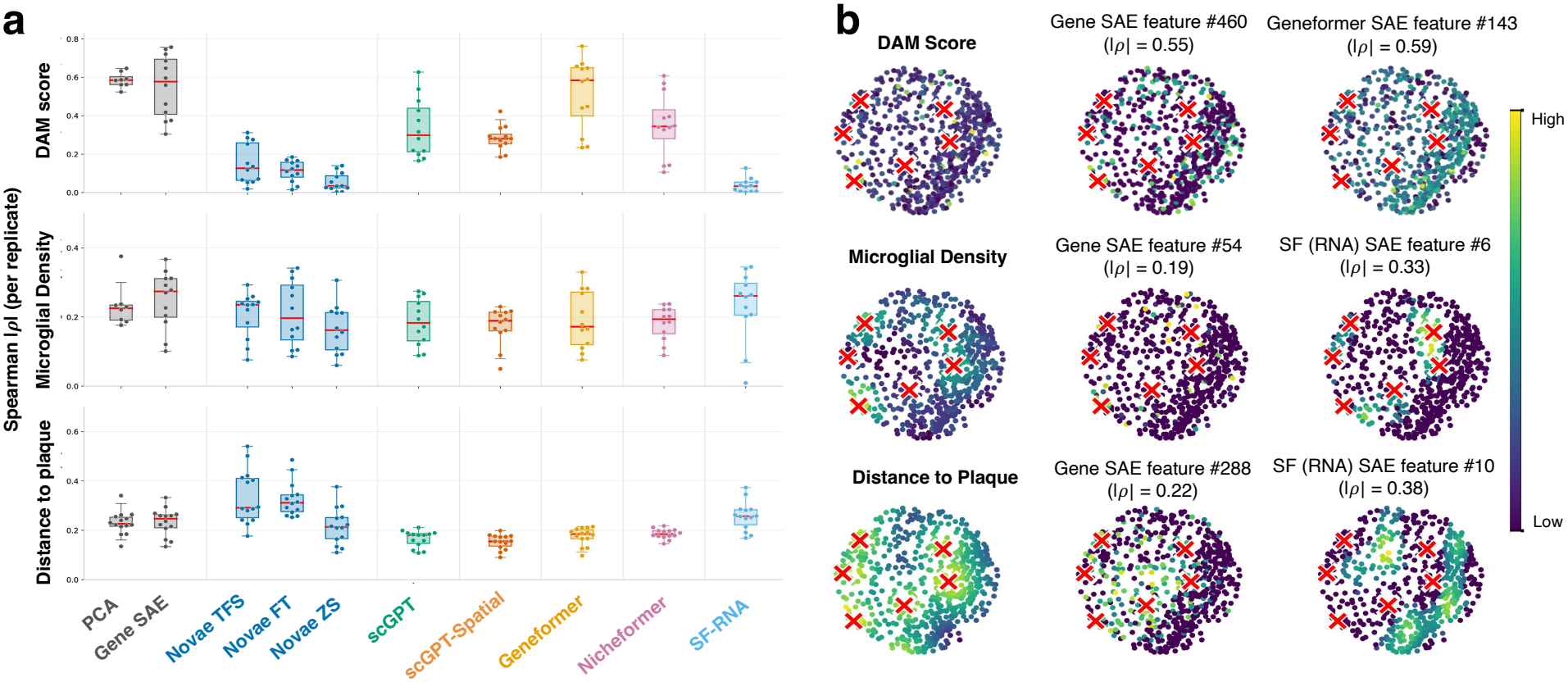
**(a)** Maximum absolute Spearman correlation |*ρ*| between plaque-associated metrics ***τ*** and SAFFRON SAE-learned features ***η*** derived from either normalized gene expression **a** or embeddings **z** of different SFM/single cell foundation models, evaluated per replicate across STARmap and MERFISH brain Alzheimer’s datasets. **(b)** Spatial zoom into a representative plaque microenvironment (MERFISH brain; n = 636 cells, 350 *µm* radius). Columns show the plaque associated metric, best Gene SAE feature (baseline), and best foundation-model SAE feature (model indicated). Red × indicates plaque centroids provided in the dataset. Absolute Spearman correlations |*ρ*| are computed within the displayed patch.

## 4 Discussion

We introduce SAFFRON, a framework for interpreting and evaluating embeddings learned by foundation models for spatial transcriptomics. SAFFRON uses a Matryoshka SAE to identify sparse, human-interpretable features from dense SFM embedding vectors, and evaluates whether the learned SAE features correlate with known local and global spatial variation in gene expression. Using SAFFRON, we find that most SFMs do not encode such multi-scale spatial variation in their embeddings: only two SFMs, Novae [28] and SpatialFusion [31], identify a continuous 1-D spatial gradient within spatial domains more accurately than non-foundation model baselines, while no SFMs identify local variation (plaque microenvironment-associated variation) better than such baselines. In particular, Novae has markedly high performance, which is likely due to explicitly modeling spatial domains in its architecture. Our findings generally suggest that current SFMs do not adequately model spatial variation in gene expression, and that future work should focus on explicitly modeling both local and global spatial variation in SFM architectures and/or pre-training objectives.

## Appendix

## A SAE ablation

We performed a systematic SAE ablation on the STARmap brain dataset (Table 1) to select the default SAE configuration used in SAFFRON. We compared Vanilla and Matryoshka SAE architectures under four sparsity modes: TopK, BatchTopK, KL, and L1. For the TopK and BatchTopK models, we swept dictionary size *M* ∈ {4*D*, 8*D*, 16*D*} and sparsity level *K/M* ∈ {5%, 10%, 15%, 20%}, where *D* is the foundation model embedding dimension. For Matryoshka SAEs, we additionally compared different scale-weighting schemes.

We evaluated each configuration using six complementary metrics: (1) Reconstruction *R*^2^ measures how accurately the SAE reconstructs the original SFM embedding. (2) Dead-latent fraction measures the proportion of dictionary elements that are never activated. (3) Cell-type probing accuracy measures whether SAE features preserve biological information relevant to cell-type classification. (4) Gini selectivity quantifies how selectively individual SAE features activate across cells. (5) Moran’s *I* measures spatial autocorrelation of SAE feature activations. (6) Feature absorption measures whether fine-grained signals are collapsed into broader latents; lower absorption indicates better feature separation.

Across the dictionary-size sweep, Novae maintained high reconstruction quality, while reconstruction performance declined at larger dictionary sizes for scGPT-Spatial and Geneformer (Figure S5). Matryoshka SAEs exhibited near-zero dead-latent fractions, while cell-type probing accuracy did not consistently improve with larger dictionaries and feature absorption generally increased. TopK and BatchTopK performed near-identically across all metrics. In contrast, KL and L1 were sensitive to the regularization strength, producing high dead-latent fractions (up to 92% and 83%, respectively) or poor reconstruction, particularly for higher-dimensional embeddings. These issues were not resolved by Matryoshka training. Equal and proportional Matryoshka weights performed comparably, with equal weights marginally better. Based on these results, we adopted BatchTopK as the sparsity constraint and selected a Matryoshka SAE for all subsequent experiments, using model-specific dictionary sizes detailed below.

## B FM evaluation details

We train a separate SAE for each foundation model embedding. For 10-dimensional embeddings (SpatialFusion), we use *M* = 40 with *K* = 4; for 64-dimensional embeddings (Novae), we use *M* = 256 with *K* = 16; and for 512- and 768-dimensional embeddings (all other models), we use *M* = 1,024 with *K* = 64. This corresponds to an active fraction of *K/M* = 10% for SpatialFusion and *K/M* = 6.25% for the remaining models. For higher-dimensional embeddings, we cap the dictionary at *M* = 1,024, as larger dictionaries did not consistently improve performance in the SAE ablation (Figure S5). Matryoshka scales are [*M/*4, *M/*2, *M*], with equal weights on the nested reconstruction losses. All SAEs are trained for 100 epochs (Adam, learning rate = 10^−3^, batch size 512), with dead-neuron resampling every 10 epochs and additive input noise annealed from *σ* = 0.02 to 0 over training.

**Table S1:** Foundation model embeddings evaluated in this study.

| **Foundation Model** | **Architecture** | **Spatial context** | **Pretraining objective** | **Dimension** |
|---|---|---|---|---|
| Geneformer [52] | BERT transformer | None | Masked gene identity prediction | 768 |
| scGPT [53] | GPT transformer | None | Masked expression value prediction | 512 |
| scGPT-Spatial [29] | Transformer + MoE decoder | Spatial patch sampling | Gene expression prediction | 512 |
| Nicheformer [30] | Transformer encoder | Cellular neighborhood transcriptomes | Masked token prediction | 512 |
| Novae [28] | Graph neural network | k-NN neighborhood graph | Self-supervised graph learning | 64 |
| SpatialFusion [31] | Transformer + graph convolutional network | k-NN graph | Multimodal reconstruction | 10 |

## C Foundation Model Fine-tuning Details

We evaluate all SFMs using their pretrained (zero-shot) embeddings and additionally fine-tune Novae and SpatialFusion using the GitHub tutorials for unsupervised fine-tuning. For SpatialFusion, the official finetuning tutorial only provides a procedure for the combined H&E and RNA modality, so we fine-tune only SF-Both. We evaluate Nicheformer and scGPT-Spatial in a zero-shot manner, as their official implementations do not provide unsupervised fine-tuning procedures. Similarly, we use pretrained embeddings for the single-cell foundation models, Geneformer and scGPT, as their official tutorials do not provide fine-tuning procedures for the spatial tasks considered here.

We followed the official fine-tuning procedures for Novae (v1.0.2) and SpatialFusion (v0.3.0). For Novae, we initialized fine-tuning using the published pretrained weights, with a maximum of 50 epochs and built-in early stopping based on training loss (minimum improvement of 0.01, patience of 5 epochs). To assess overfitting, we trained and fine-tuned Novae on the MERFISH Alzheimer’s disease dataset (12 mice) and evaluated the training objective on held-out mice (4 mice) after each epoch, across three random seeds. The held-out loss decreased alongside the training loss and plateaued without increasing, suggesting no clear evidence of overfitting (Figure S6a). For SpatialFusion, we followed the official fine-tuning procedure on datasets with paired H&E and RNA. Since the training loss continued to decrease or stabilize, we fine-tuned the paired autoencoder and spatial graph convolutional model for 15 epochs each, at the upper end of the recommended 5–15 epoch range (Figure S6a).

### Cross-dataset generalization

To assess cross-dataset generalization, we trained Novae from scratch on the mouse kidney dataset and evaluated it on the human colorectal tumor dataset without further training. The mean maximum absolute Spearman correlation across spatial domains was 0.41, lower than that of models trained or fine-tuned on colorectal (0.66–0.72), but substantially above the permutation-based null (Figure S6b). This suggests that some spatial gradient information remains recoverable across datasets, although recovery is weaker than with training on the target dataset.

**Figure S1:**
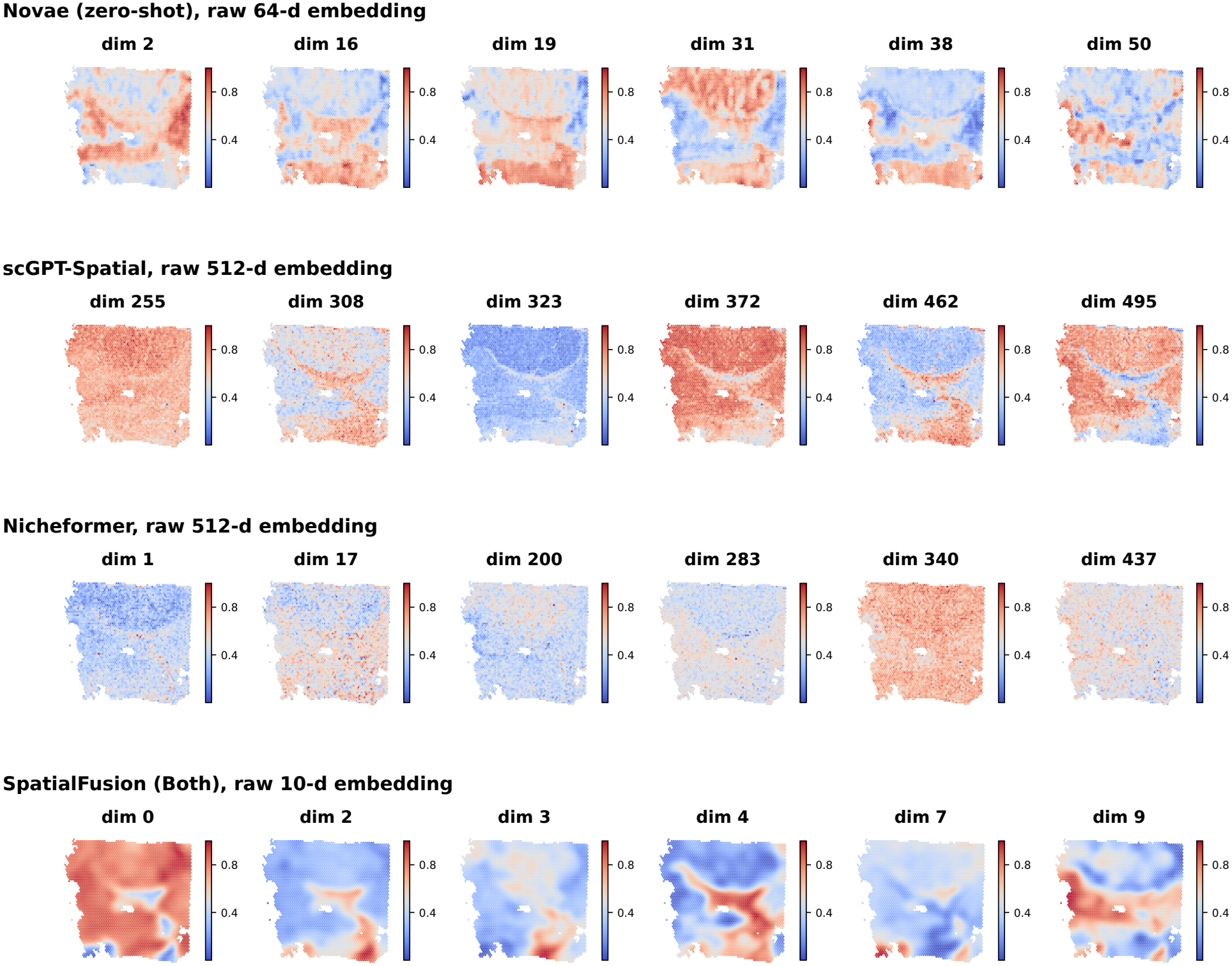
Spatial visualization of raw embedding dimensions **z** from Novae (zero-shot, 64-dimensional), scGPT-Spatial (512-dimensional), Nicheformer (512-dimensional), and SpatialFusion (Both, 10-dimensional) on the colorectal tumor dataset. Each spatial map shows the normalized values of an individual embedding dimension across the tissue section, without SAE decomposition.

**Figure S2:**
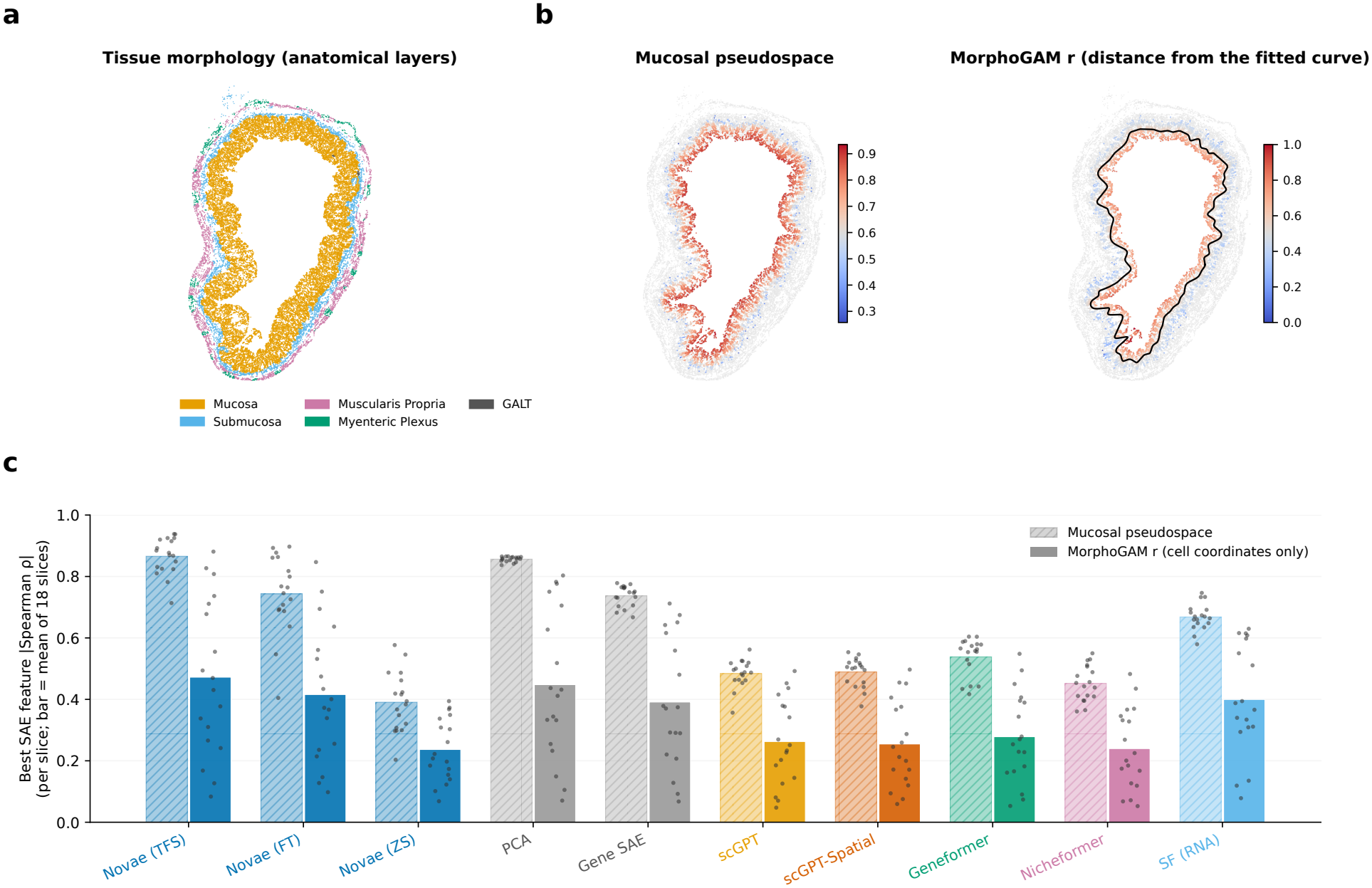
**(a)** Representative GI tract tissue section, colored by anatomical layer annotations provided in the dataset. **(b)** MorphoGAM [55] radial coordinate *r* for the same section, calculated as the signed distance of each cell from a curve (black) fitted through the spatial positions of mucosal enterocytes, independently of gene expression. **(c)** Maximum absolute Spearman correlation |*ρ*| between SAE-learned features ***η*** derived from normalized gene expression **a** or embeddings **z** of different foundation models, as well as PCA features, and either the mucosal pseudospace (hatched) or MorphoGAM *r* (solid), evaluated across 18 GI tract sections. Bars indicate mean correlations across sections, and dots represent individual sections.

**Figure S3:**
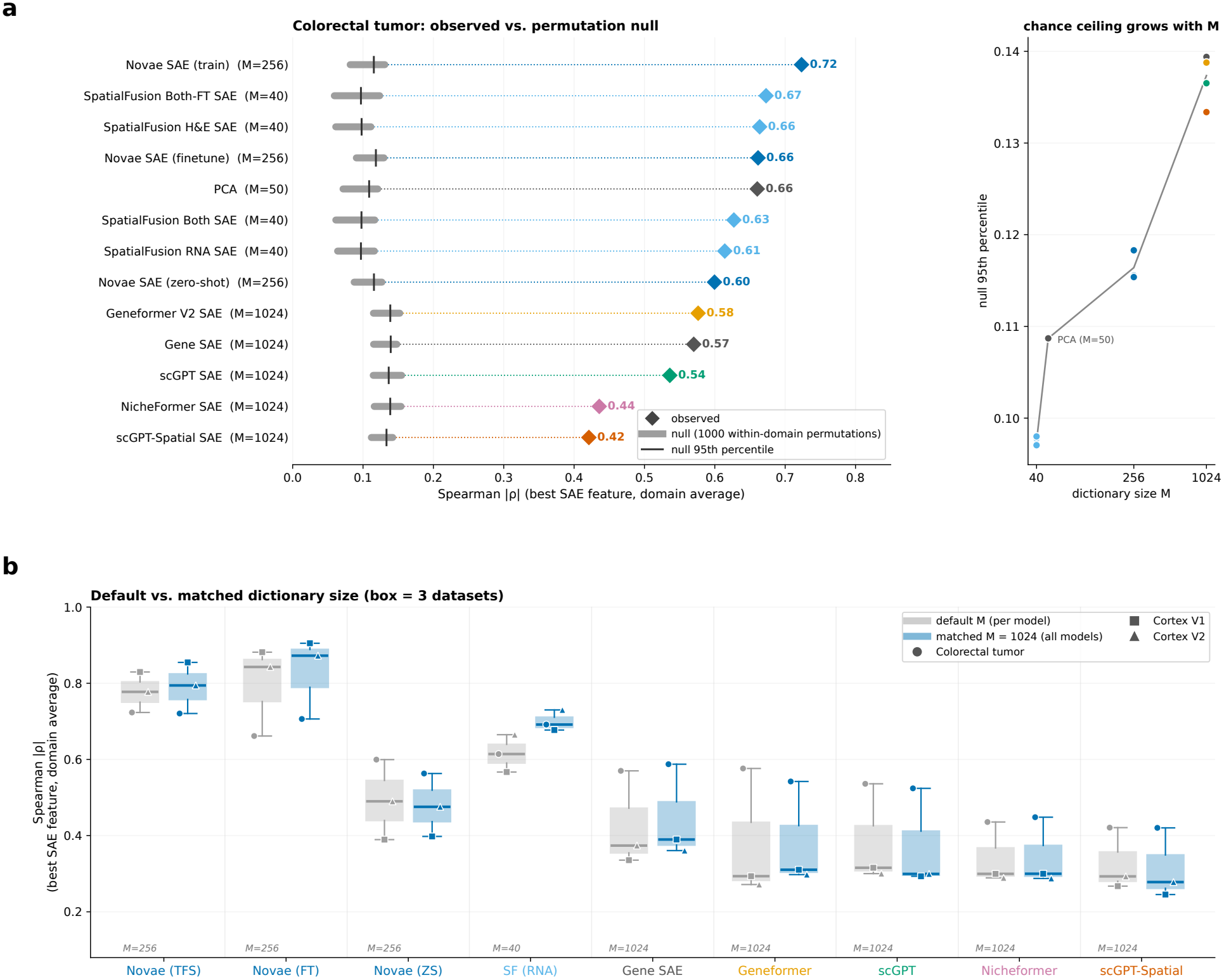
**(a)** Permutation-based null analysis of spatial gradient recovery in the colorectal tumor dataset. Left: Observed gradient recovery (maximum absolute Spearman correlation |*ρ*| across SAE features) averaged across spatial domains, compared against null distributions obtained by randomly permuting the 1-D spatial coordinate 1,000 times within each domain. Gray distributions show the permutation nulls, with the 95th percentile indicated by vertical lines. Right: 95th percentile of the null distribution as a function of dictionary size *M*. **(b)** Comparison of gradient recovery using each model’s original SAE dictionary size and a matched dictionary size (*M* = 1024) across three datasets (colorectal tumor, cortex V1, and cortex V2). Gray and blue boxplots indicate the original and matched dictionary sizes, respectively, with markers representing individual datasets.

**Figure S4:**
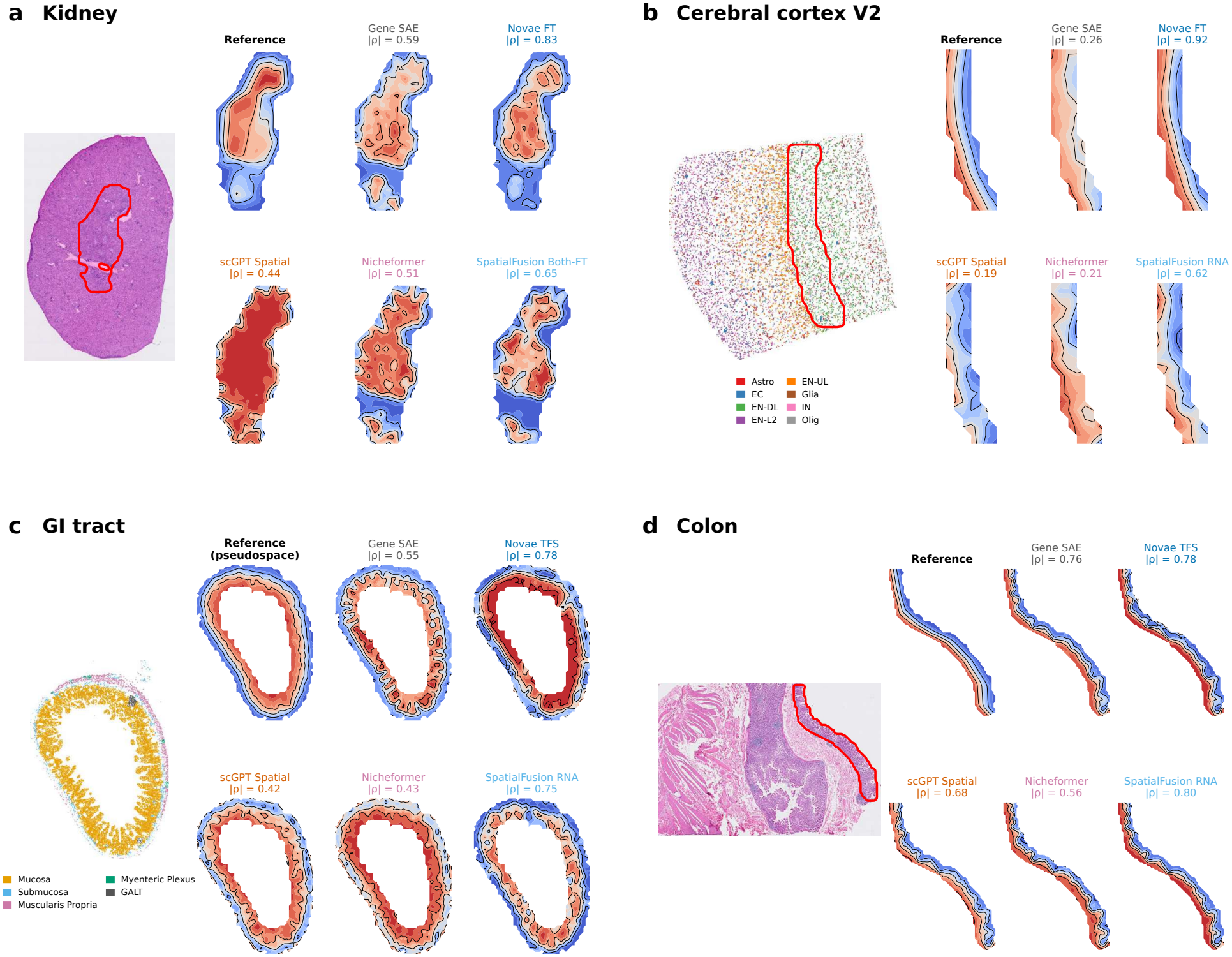
**(a)–(d)** (Left) H&E images for kidney (a) and colon (d), cell type annotations for cerebral cortex V2 (b), and anatomical layer annotations for GI tract (c). Outlined regions denote the spatial domains visualized on the right. (Right) Reference 1-D spatial axis ***τ*** and SAFFRON SAE-learned features with the highest absolute Spearman correlation to ***τ*** for each model, with |*ρ*| indicated.

**Figure S5:**
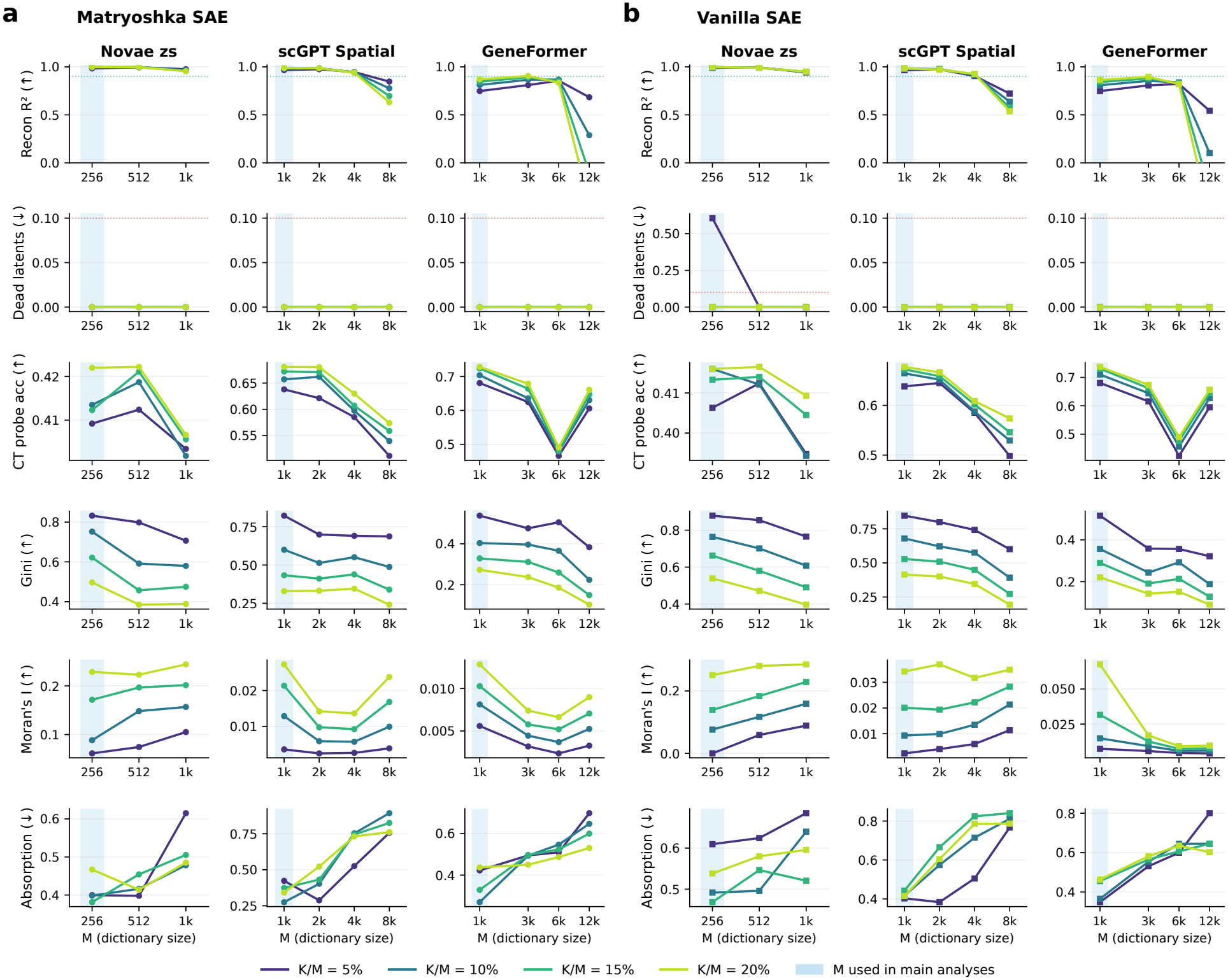
Comparison of **(a)** Matryoshka and **(b)** Vanilla SAEs trained on Novae (zero-shot), scGPT-Spatial, and Geneformer embeddings from the STARmap brain dataset. Rows show reconstruction *R*^2^, dead-latent fraction, cell-type probing accuracy, Gini selectivity, Moran’s *I*, and feature absorption across dictionary sizes *M*. Colored lines represent sparsity levels *K/M* ∈ {5%, 10%, 15%, 20%}. Blue shading indicates the dictionary size used in the main analyses.

**Figure S6:**
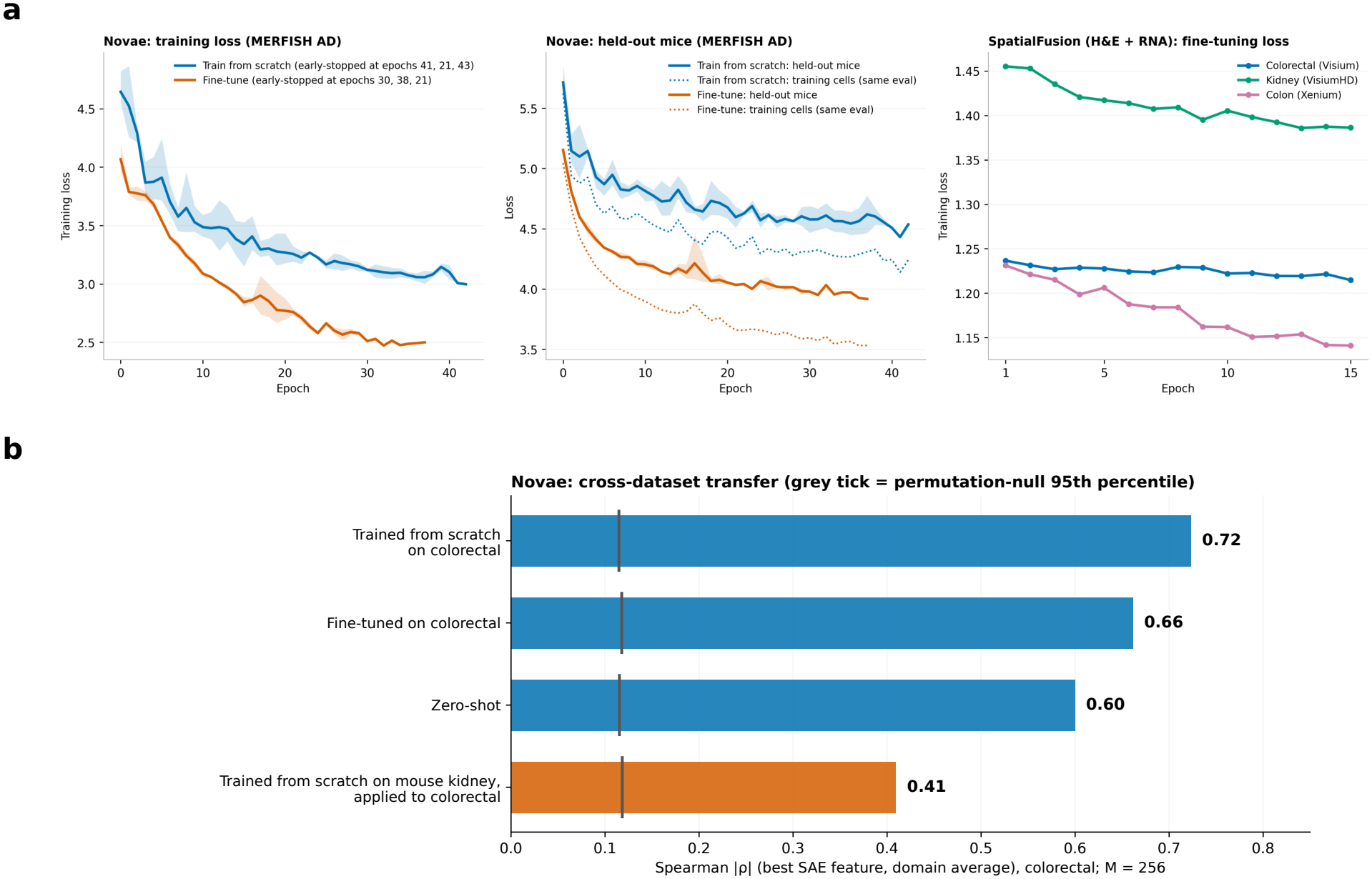
**(a)** Training and fine-tuning loss curves for Novae and SpatialFusion. Left: Novae training loss on the MERFISH Alzheimer’s disease dataset, trained from scratch or fine-tuned from pretrained weights. Lines indicate the mean across three random seeds, with shaded regions showing the range. Middle: Corresponding loss on held-out mice (solid) and training cells (dotted). Right: SpatialFusion fine-tuning loss on three datasets (colorectal, kidney, colon) with paired H&E and RNA over 15 epochs. **(b)** Cross-dataset evaluation of Novae trained from scratch on mouse kidney and applied to the human colorectal tumor dataset without further training, compared with Novae trained from scratch, fine-tuned, or evaluated zero-shot on colorectal. Bars indicate the mean across spatial domains of the maximum absolute Spearman correlation |*ρ*| across SAE features. Gray lines indicate the 95th percentile of the permutation-based null distribution.

